# Loss of E2F7 confers resistance to poly-ADP-ribose polymerase (PARP) inhibitors in BRCA2-deficient cells

**DOI:** 10.1101/294371

**Authors:** Kristen E. Clements, Tanay Thakar, Claudia M. Nicolae, Xinwen Liang, Hong-Gang Wang, George-Lucian Moldovan

## Abstract

BRCA proteins are essential for Homologous Recombination DNA repair, and their germline or somatic inactivation is frequently observed in human tumors. Understanding the molecular mechanisms underlying the response to chemotherapy of BRCA-deficient tumors is paramount for developing improved personalized cancer therapies. While PARP inhibitors have been recently approved for treatment of BRCA-mutant breast and ovarian cancers, resistance to these novel drugs remains a major clinical problem. Several mechanisms of chemoresistance in BRCA2-deficient cells have been identified. Rather than restoring normal recombination, these mechanisms result in stabilization of stalled replication forks, which normally are subjected to degradation in BRCA2-mutated cells. Here, we show that the transcriptional repressor E2F7 controls chemoresistance in BRCA2-deficient cells. We found that E2F7 depletion restores PARP inhibitor and cisplatin resistance in BRCA2-depleted cells. Moreover, we show that the mechanism underlying this activity involves increased expression of RAD51, a target for E2F7-mediated transcriptional repression, which enhances both Homologous Recombination DNA repair, and replication fork stability in BRCA2-deficient cells. Our work describes a new mechanism of chemotherapy resistance in BRCA2-deficient cells, and identifies E2F7 as a novel biomarker for tumor response to PARP inhibitor therapy.

## INTRODUCTION

Improved precision therapy is essential for increasing life expectancy of cancer patients. PARP1 is a member of the poly-ADP-ribosyltransferase family, catalyzing formation of poly-ADP-ribose chains on target protein substrates (1). PARP1 has diverse substrates and regulates essential cellular processes including DNA replication, DNA repair, and transcription. Recently, PARP1 inhibitors have emerged as novel cancer therapeutics, based on groundbreaking work showing that PARP1 is essential for cellular viability in cells with compromised Homologous Recombination DNA repair (HR) (2-4). Inability to perform PARP1-mediated repair of single stranded DNA breaks leads to replication fork collapse and double strand break formation. In the absence of efficient HR, this results in cell death, underlying the synthetic lethality interactions between PARP1 and HR genes. HR deficiency conferred by germline or somatic mutations in BRCA1, BRCA2, RAD51C, Fanconi Anemia genes and other members of the pathway is observed in a large proportion of adult cancers, including breast, ovarian, pancreatic, prostate and others (5,6). Several PARP inhibitors (PARPi) (olaparib, rucaparib, and niraparib) have been FDA-approved for single agent treatment of HR-deficient ovarian and breast cancers.

More recently, it was shown that PARPi can also be effective against HR-proficient tumors, through a newly described activity known as PARP-trapping-which results in crosslinking of the PARP1 protein to DNA (7). These DNA-protein crosslinks can block DNA replication and transcription. Through this mechanism, PARPi act as efficient chemo- and radio-sensitizers (8-10). Thus, use of PARPi is likely to be significantly extended in the near futures to many different cancers, regardless of HR (BRCA) status. Indeed, there are currently more than 20 active clinical trials involving PARPi, in tumors ranging from breast to bone to brain, in both children and adults (11).

BRCA2 is an essential HR protein, which catalyzes the loading of RAD51 molecules onto resected DNA at double strand breaks (12). RAD51 loading is required for the subsequent strand invasion and Holliday junction formation steps of the recombination process. BRCA2 was also shown to be required for genomic stability under replication stress conditions (13,14). Upon replication fork stalling at sites of DNA lesions, potentially including trapped PARP1, a set of DNA translocases including ZRANB3, HLTF and SMARCAL1 reverse the fork by annealing the nascent strands of the two newly formed chromatids. The resulting double stranded DNA break (DSB) end needs to be stabilized by BRCA2-mediated loading of RAD51, which protects it against degradation by the MRE11 nuclease (15-17).

While PARPi have excellent anti-tumor activity, they often show only limited efficacy in the clinic. For example, even though olaparib treatment tripled 12-month progression free survival in BRCA2 deficient patients, still only 65% of the olaparib-treated group reached this milestone, indicating that resistance is an important clinical problem (18). Mechanisms of acquired resistance include genetic reversion of BRCA1 and BRCA2 mutations, as well as rewiring of the DNA damage response to restore HR in BRCA1-deficient cells by suppressing recombination-inhibitory activities such as 53BP1 or RIF1 (19-22). In contrast, in BRCA2-deficient cells, known mechanisms of resistance do not restore HR, but instead act by protecting stalled replication forks against nucleolytic degradation (20,23).

E2F7 is a member of the E2F transcription factor family. Together with E2F8, they are considered atypical E2F family members as they mediate transcription repression rather than activation (24,25).

E2F7 levels are induced by DNA damage (26). E2F7 transcriptional repression targets include replication proteins such as CDC6 and MCM2-thereby its induction by DNA damage contributes to G1/S-arrest (27,28). However, among its targets for repression are also HR proteins, including RAD51 and BRCA1 (28). Here, we show that E2F7 promotes sensitivity to PARPi, and its loss can rescue chemosensitivity of BRCA2deficient cells by promoting both HR and fork stability.

## MATERIAL AND METHODS

### Cell culture and protein techniques

Human HeLa, HCC1395, 293T, and U2OS cells were grown in DMEM, while SH-SY5Y were grown in DMEM/F12 (1:1). Media was supplemented with 10% Fetal Calf Serum. For BRCA2 gene knockout, the commercially available BRCA2 CRISPR/Cas9 KO plasmid was used (Santa Cruz Biotechnology sc-400700). Single transfected cells were FACS-sorted into 96-well plates using a BD FACSAria II instrument. Resulting colonies were screened by Western blot. Cell extracts, chromatin fractionation, and Western blot experiments were performed as previously described (29-31). Antibodies used for Western blot are: BRCA2 (Bethyl A303-434A), GAPDH (Santa Cruz Biotechnology sc-47724), RAD51 (Santa Cruz Biotechnology sc-8349), Vinculin (Santa Cruz Biotechnology sc-73614). For gene knockdown, cells were transfected with Stealth siRNA (Life Tech) using Lipofectamine RNAiMAX reagent. The siRNA targeting sequences used are: E2F7 #1: GGACGATGCATTTACAGATTCTCTA; E2F7 #2: GACTATGGGTAACAGGGCATCTATA; E2F7 #3: AAACAAAGGTACGACGCCTCTATGA (used for E2F7 knockdown unless otherwise mentioned); BRCA2: GAGAGGCCTGTAAAGACCTTGAATT; RAD51: CCATACTGTGGAGGCTGTTGCCTAT.

### Functional assays

To measure drug sensitivity, cells were seeded in 96-well plates and incubated with indicated drug concentrations for 3 days. Cellular viability was assayed using the CellTiterGlo reagent (Promega) according to the manufacturer’s instructions. Apoptosis Annexin V measurements were performed using the FITC Annexin V kit (Biolegend 640906) according to manufacturer’s instructions, using a BD FACSCanto 10 flow cytometer. DR-GFP assays were performed as previously described (32).

### Quantification of gene expression by real-time quantitative PCR (RT-qPCR)

Total mRNA was purified using TRIzol reagent (Life Tech) according to the manufacturer’s instructions. To generate cDNA, 1 μg RNA was subjected to reverse transcription using the RevertAid Reverse Transcriptase Kit (Thermo Fisher Scientific) with oligo dT primers. Real-time qPCR was performed with PerfeCTa SYBR Green SuperMix (Quanta), using a CFX Connect Real-Time Cycler (BioRad). The cDNA of GAPDH gene was used for normalization. Primers used were: E2F7 for: GGAAAGGCAACAGCAAACTCT; E2F7 rev: TGGGAGAGCACCAAGAGTAGAAGA; RAD51 for: TGCTTATTGTAGACAGTGCCACC; RAD51 rev: CACCAAACTCATCAGCGAGTC; GAPDH for: AAATCAAGTGGGGCGATGCTG; GAPDH rev: GCAGAGATGATGACCCTTTTG

### DNA Fiber Assay

293T cells, pretreated with the indicated siRNA oligonucleotides, were incubated with 100μM IdU for 30 minutes. Cells were washed with PBS and incubated with 4mM HU (with or without 50μM Mirin as indicated) for 4h. Following removal of HU media and a PBS wash, fresh media containing 100μM CldU was added for another 30 minutes. Next, cells were harvested and DNA fibers were obtained using the FiberPrep kit (Genomic Vision) according to the manufacturer’s instructions. DNA fibers were stretched on glass slides using the FiberComb Molecular Combing instrument (Genomic Vision). Slides were incubated with primary antibodies (Abcam 6326 for detecting CIdU; BD 347580 for detecting IdU; Millipore Sigma MAB3034 for detecting DNA), washed with PBS, and incubated with Cy3, Cy5, or BV480-coupled secondary antibodies (Abcam 6946, Abcam 6565, BD Biosciences 564879). Following mounting, slides were imaged using a Leica SP5 confocal microscope. At least 200 tracts were quantified for each sample.

## RESULTS

### Loss of E2F7 reverses the PARPi sensitivity of BRCA2deficient cells

In order to investigate mechanisms of PARPi resistance in BRCA-deficient cells, we first created a BRCA2-knockout (labeled BRCA2^KO^) HeLa cell line using the CRISPR/Cas9 technology. BRCA2^KO^ cells lack any detectable BRCA2 protein expression by Western blot (Fig. 1A) and have similar sensitivity to olaparib as parental cells treated with BRCA2-targeting siRNA (Fig. 1B). In order to investigate its involvement in mediating PARPi-sensitivity in BRCA2-deficient cells, we knocked down E2F7 in BRCA2^KO^ cells and measured olaparib sensitivity. E2F7 knockdown resulted in significant rescue of olaparib sensitivity of these cells (Fig. 1C). In order to rule out a non-specific effect caused by the CRISPR editing, we also knocked-down E2F7 in parental HeLa cells at the same time with BRCA2 depletion. E2F7 knockdown with three different siRNA oligonucleotides could rescue olaparib sensitivity of BRCA2-knockdown cells (Fig. 1D). All three siRNA oligonucleotides can efficiently knock-down E2F7 expression as shown by qPCR-based detection of E2F7 mRNA levels (Fig. 1E). Further confirming the specificity of the phenotype, Western blot experiments showed that BRCA2 is equally depleted in cells treated with BRCA2 siRNA alone or in combination with E2F7 siRNA (Fig. 1F). Olaparib treatment is known to induce apoptosis in BRCA2-deficient cells (33). E2F7 depletion did not only rescue cellular viability of olaparib-treated BRCA2-deficient cells, but also olaparib-induced apoptosis (Fig. 1G). Altogether, these results indicate that loss of E2F7 is a novel mechanism of PARPi resistance in BRCA2-deficient cells.

**Figure 1.**
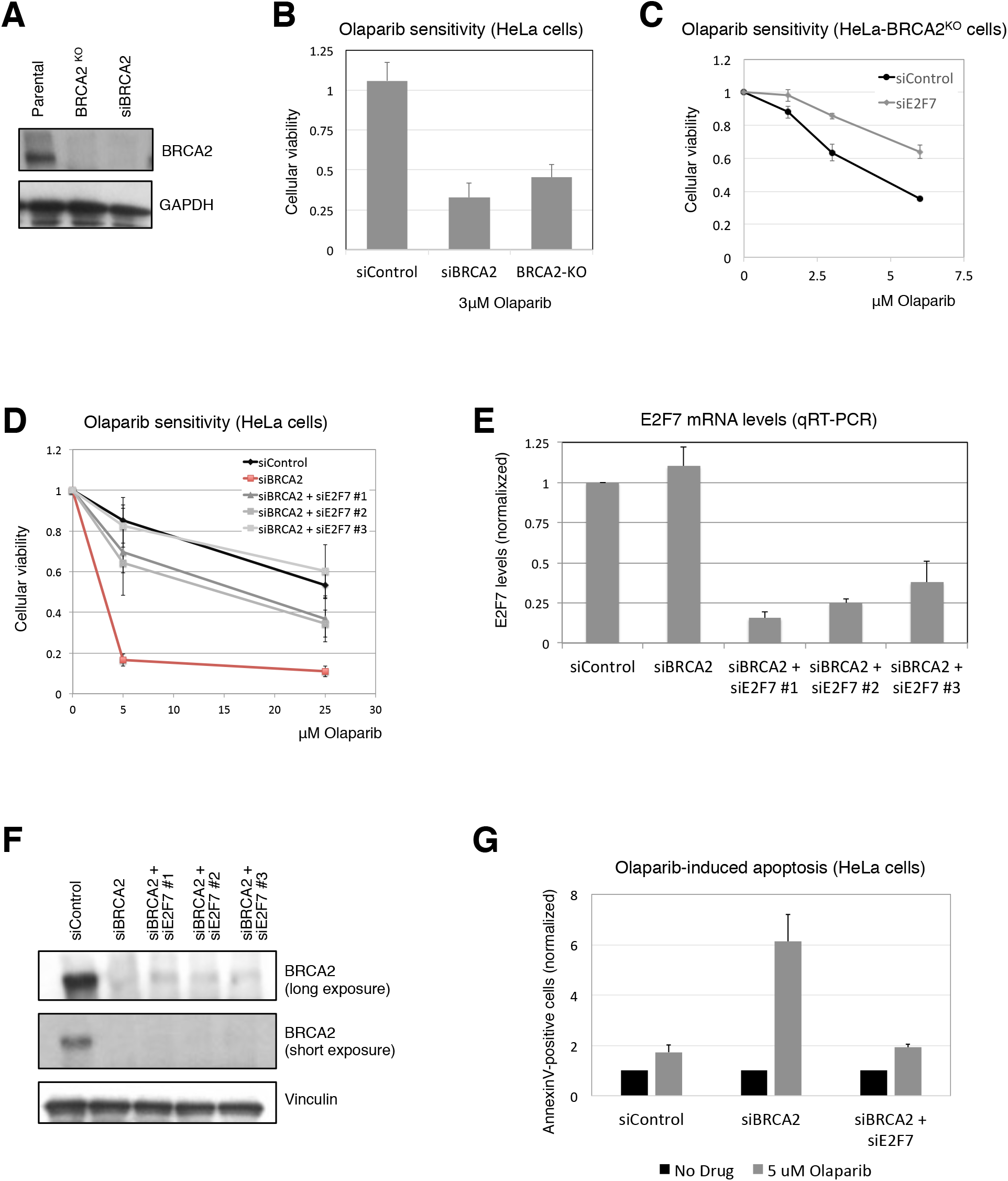
E2F7 depletion in BRCA2-deficient HeLa cells results in olaparib resistance. **A**. Western blot showing loss of BRCA2 expression in HeLa cells with CRISPR/Cas9-mediated BRCA2 knockout. **B**. BRCA2^KO^ HeLa cells have similar olaparib sensitivity as HeLa cells treated with BRCA2 siRNA. Results are shown as normalized to control (no drug treatment) for each sample. The average of three experiments, with error bars as standard deviations, is shown. **C, D**. E2F7 knockdown rescues the olaparib sensitivity of BRCA2-knockout (**C**) and BRCA2-knockdown (**D**) HeLa cells. The average of three experiments, with error bars as standard deviations, is shown. **E**. Quantitative RT-PCR experiment showing efficient E2F7 knockdown by the siRNA oligonucleotides employed. HeLa cells were treated with the indicated siRNA oligonucleotides, than incubated with 10μM olaparib for 24h before harvesting. The average of three experiments, with error bars as standard deviations, is shown. **F**. Western blot showing that BRCA2 is efficiently knocked down by the siRNA oligonucleotide employed singly or in combination with E2F7 siRNA oligonucleotides. HeLa cells were treated with the indicated siRNA oligonucleotides, than incubated with 10μM olaparib for 24h before harvesting. **G**. Quantification of AnnexinV-positive cells indicating that E2F7 knockdown suppresses olaparib-induced apoptosis of BRCA2-depleted cells. HeLa cells were treated with the indicated siRNA oligonucleotides, than incubated with 5μM olaparib for 3 days. Results are presented as normalized to control (no drug treatment condition) for each knockdown sample. The average of three independent experiments, with standard deviations as error bars, is shown.

We next investigated if the suppression of olaparib sensitivity by loss of E2F7 is restricted to HeLa cells, or is a general phenomenon. BRCA2 depletion in HCC1395 breast cancer cells resulted in olaparib sensitivity, which was rescued by E2F7 knockdown (Fig. 2A). Similarly, olaparib sensitivity of BRCA2-depleted SH-SY5Y neuroblastoma cells was rescued by E2F7 depletion (Fig. 2B). Preclinical studies showed that PARP inhibitors display synergistic effects with Camptothecin in BRCA-proficient neuroblastoma cell lines, which represents the basis for current clinical trials combining PARPi and genotoxic chemotherapy in neuroblastoma treatment (34). E2F7 knockdown suppressed the sensitivity of SH-SY5Y cells to olaparib-camptothecin combination treatment (Fig. 2C), indicating that the new PARPi resistance mechanism we identified is relevant for other putative clinical applications of PARPi treatment, namely combination therapy in BRCA-proficient cells.

**Figure 2.**
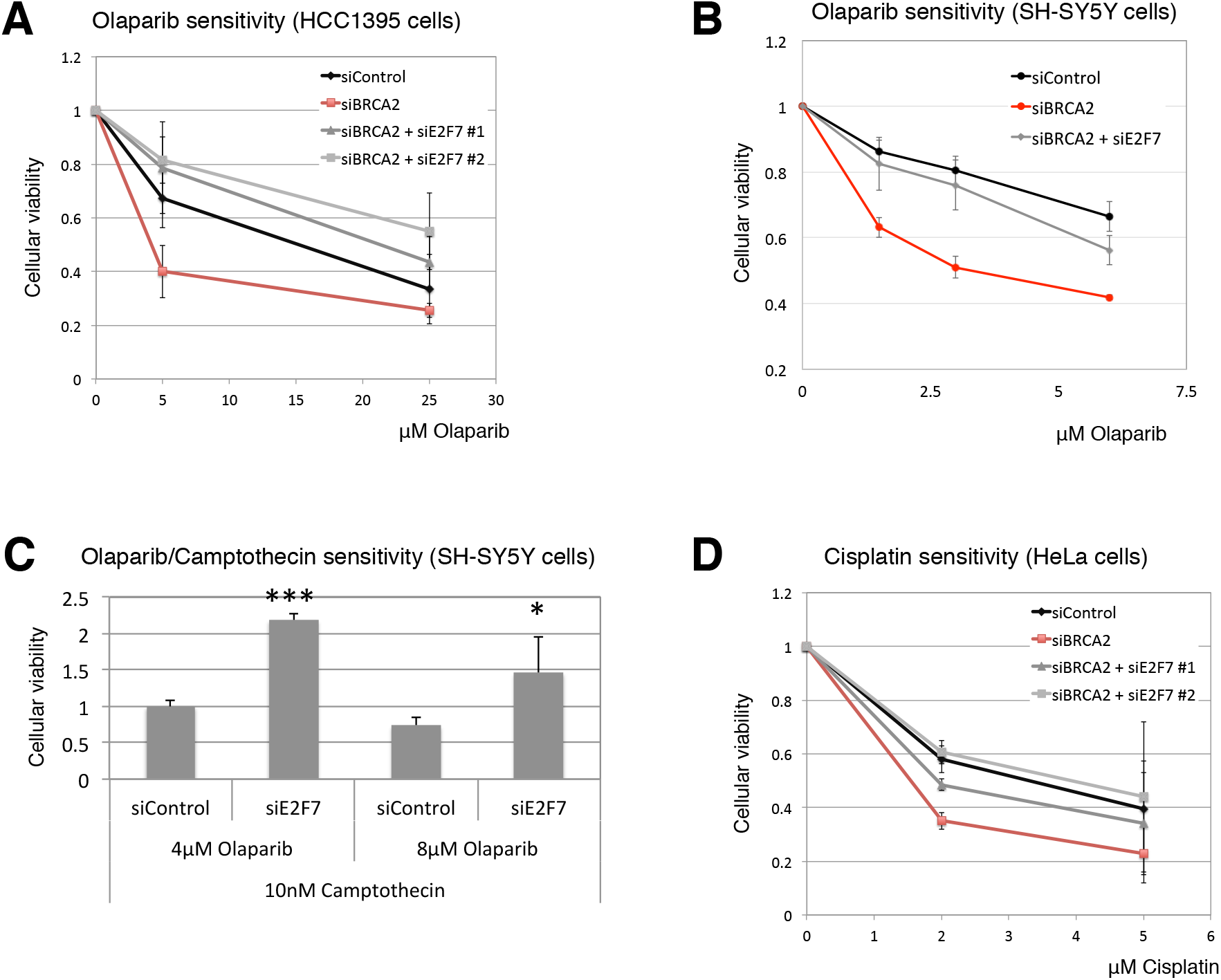
Loss of E2F7 suppresses BRCA2-induced chemosensitivity. **A, B**. E2F7 knockdown rescues the olaparib sensitivity of BRCA2-knockdown HCC1395 breast cancer (**A**) and SH-SY5Y neuroblastoma (**B**) cells. HCC1395 cells are BRCA1-mutant, which accounts for their increased olaparib sensitivity. The average of three experiments, with error bars as standard deviations, is shown. **C**. E2F7 depletion increases resistance of SH-SY5Y cells to olaparib-camptothecin combination treatment. The average of three technical replicates, with error bars as standard deviations, is shown. **D**. E2F7 knockdown rescues the cisplatin sensitivity of BRCA2-knockdown HeLa cells. The average of three experiments, with error bars as standard deviations, is shown.

BRCA2 deficient cells are sensitive to many other genotoxic agents used in cancer therapy, including cisplatin (35). In order to test if this E2F7-mediated response is specific to PARPi, we treated E2F7/BRCA2-co-depleted HeLa cells with cisplatin. E2F7 knockdown was able to rescue the cisplatin sensitivity of BRCA2-depleted cells (Fig. 2D), indicating that E2F7 has a broad impact on the chemosensitivity of BRCA2-deficient cells. Altogether, these findings show that E2F7 is a novel regulator of DNA damage sensitivity of BRCA2-deficient cells.

### Loss of E2F7 restores RAD51-mediated Homologous Recombination in BRCA-deficient cells

Previously, RAD51 was identified as a possible target for transcriptional repression by E2F7 (28). RAD51 is an essential HR factor, which is loaded by BRCA2 on resected ssDNA and catalyzes the strand invasion step in the recombination process (12). Thus, we decided to investigate the levels of RAD51 in BRCA2-depleted HeLa cells treated with Olparaib. E2F7 knockdown resulted in increased RAD51 expression, at both the mRNA and protein levels (Fig. 3A, B). Moreover, E2F7 knockdown resulted in an increase in the levels of chromatin bound RAD51 upon olaparib treatment (Fig. 3C). These results suggest that E2F7 depletion may promote olaparib resistance in BRCA2-deficient cells by increasing RAD51 levels.

**Figure 3.**
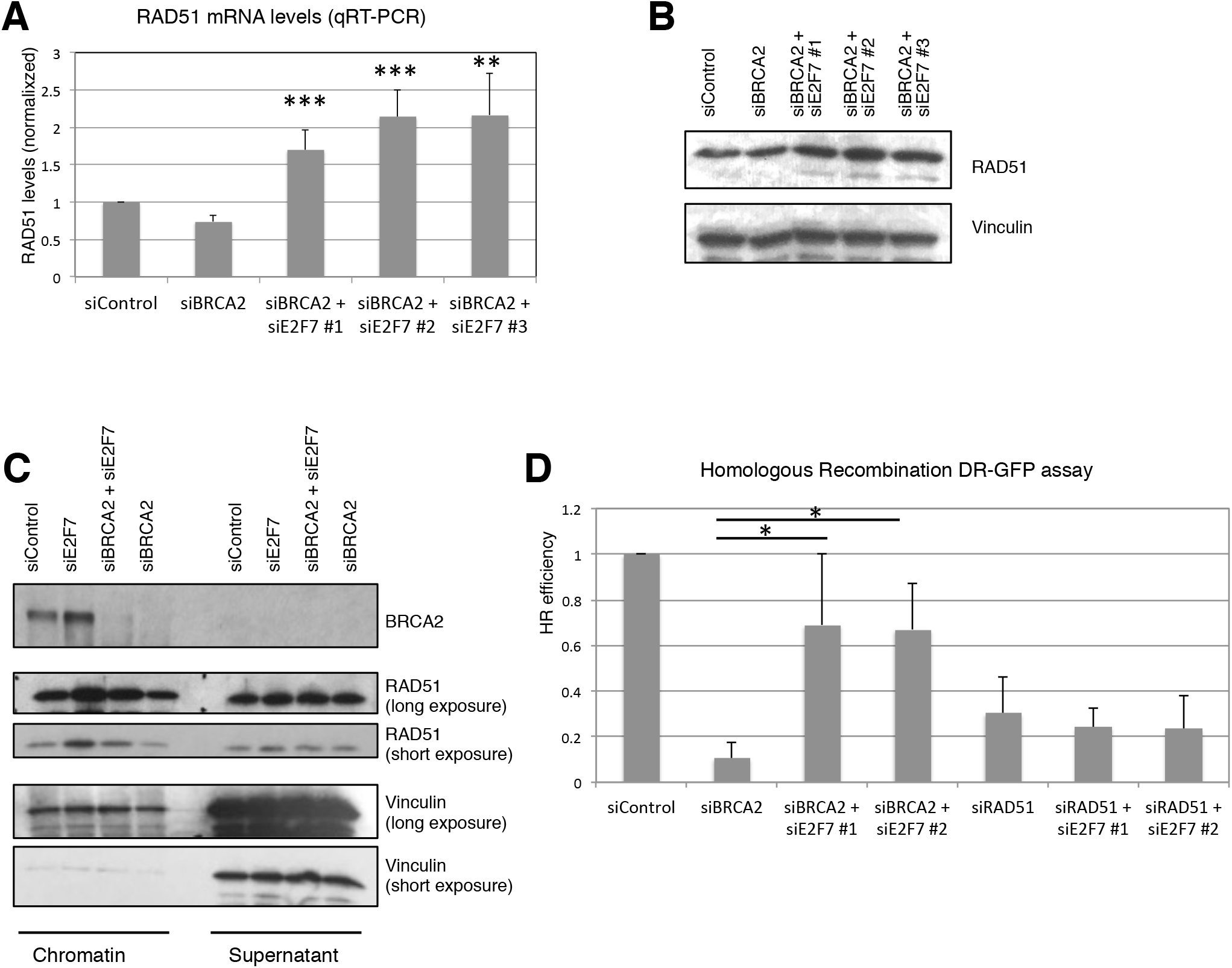
E2F7 knockdown increases RAD51 expression and HR levels in BRCA2-deficient cells. **A**. Quantitative RT-PCR experiment showing increased RAD51 mRNA levels upon E2F7 depletion in BRCA2-knockdown HeLa cells incubated with 10μM olaparib for 24h before harvesting. The average of three experiments, with error bars as standard deviations, is shown. Asterisks denote statistical significance compared to siBRCA2 condition (using the two-tailed equal variance TTEST). **B**. Western blot showing that E2F7 knockdown results in increased RAD51 protein levels in HeLa cells treated with 10μM olaparib for 24h. **C**. Chromatin fractionation experiment showing that E2F7 depletion results in increased chromatin-bound RAD51. HeLa cells were treated with 10μM olaparib for 24h. Vinculin is used as loading control. **D**. DR-GFP assay in U2OS cells showing that E2F7 depletion restores HR levels in BRCA2-knockdown cells, but not in RAD51-knockdown cells. The average of three experiments, with error bars as standard deviations, is shown. Asterisks indicate statistical significance (using the two-tailed equal variance TTEST).

Next we investigated the functional outcomes of increased RAD51 levels in BRCA2-deficient cells. We reasoned that increased RAD51 loading on damaged DNA may partly alleviate the impact of BRCA2 loss and restore HR proficiency. To address this, we employed the DR-GFP assay that measures HR between direct repeats in U2OS cells (32). As expected, BRCA2 knockdown significantly reduced HR. However, co-depletion of E2F7 could rescue HR in BRCA2-deficient cells (Fig. 3D). In contrast, E2F7 depletion could not rescue the HR defect conferred by loss of RAD51 (Fig. 3D), indicating that the E2F7-mediated rescue observed in BRCA2-deficient cells requires RAD51 expression. Altogether, these results indicate that loss of E2F7 increases RAD51 levels, which promotes HR in BRCA-deficient cells.

### E2F7 regulates replication fork stability in BRCA2-deficient cells

Recently, degradation of stalled replication forks has emerged as a novel activity underlining chemosensitivity of BRCA2-deficient cells. In normal cells, BRCA2 protects against fork degradation by loading RAD51 on forks arrested at sites of DNA damage. In BRCA-deficient cells, RAD51 cannot be loaded thus rendering stalled forks susceptible to degradation by MRE11 nuclease (13,14,16). In order to test the impact of E2F7 on fork stability in BRCA-deficient cells, we employed the DNA fiber assay, which allows detection and quantification of nascent DNA strands at molecular level. We measured the length of replication tracts upon treatment with hydroxyurea (HU), an established model of fork degradation in BRCA-deficient cells (15,16,36). In line with previous reports (13,14,16), BRCA2 knockdown resulted in HU-induced degradation of nascent DNA, which could be suppressed by treatment with MRE11 inhibitor Mirin (Fig. 4). Importantly, E2F7 depletion could also rescue the HU-induced fork degradation phenotype of BRCA2-knockdown cells (Fig. 4B). These results indicate that increased RAD51 levels upon E2F7 depletion are enough to promote stability of stalled replication forks in BRCA2-deficient cells.

**Figure 4.**
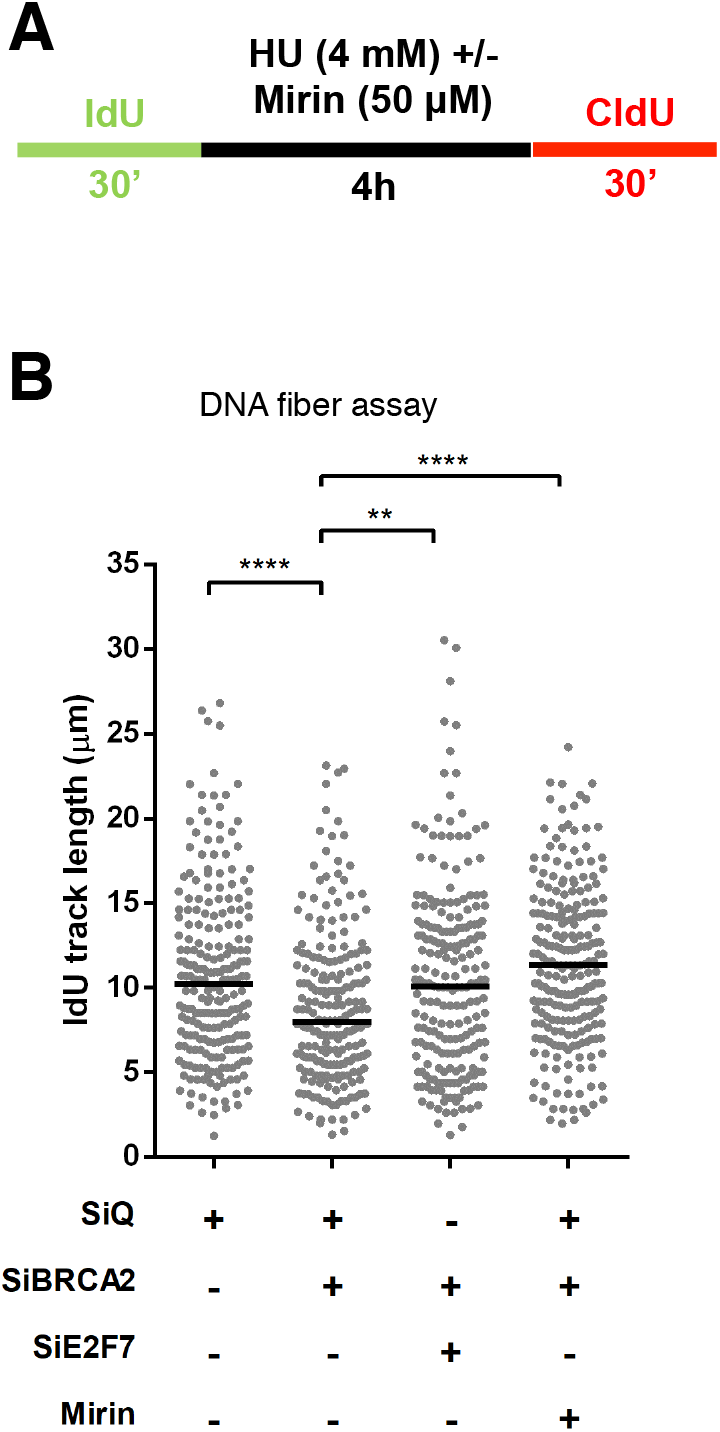
E2F7 controls replication fork stability in BRCA2-deficient cells. **A**. Schematic representation of the experimental setup for the DNA fiber combing experiment. **B**. Quantification of the IdU tract length. BRCA2 knockdown reduces the length of the IdU tract upon HU treatment, indicating that the nascent strand is degraded. This degradation can be suppressed by E2F7 depletion, or as control, by incubation with the MRE11 inhibitor mirin. The median is indicated.

## DISCUSSION

Identifying clinical biomarkers of PARPi resistance is paramount for more targeted usage of these promising drugs in cancer treatment. Our work identifies loss of E2F7 as a new mechanism of PARPi and cisplatin resistance of BRCA2-deficient cells. We show that E2F7 depletion results in increased levels of RAD51, a previously described target for transcriptional repression by E2F7. This, in turn, promotes both HR and fork stability in BRCA2-deficient cells. Moreover, we show that loss of E2F7 also promotes resistance of BRCA-proficient cancer cells to PARPi-Camptothecin combination therapy, a novel mode of employing PARPi, currently undergoing clinical trials. Thus, our work indicates that E2F7 levels may represent a critical biomarker predicting PARPi responses of human tumors. Moreover, upstream regulators of E2F7 may also serve as biomarkers and provide opportunities for therapeutic intervention. Recently, Chk1 was shown to phosphorylate E2F7 thereby restricting its activity in response to DNA damage (37). We thus predict that in BRCA2-deficient cells, Chk1 activity, by inhibiting E2F7, promotes PARPi resistance. Pharmacological inhibition of this pathway may potentially be employed to restore PARPi sensitivity in tumors with acquired resistance.

Previous studies have shown that mechanisms of PARPi resistance differ between BRCA1 and BRCA2-deficient cells. Sensitivity of BRCA1-deficient cells can be suppressed by restoring HR, while in BRCA2-deficient cells resistance occurs through promoting replication fork stability and protection against nucleolytic degradation of stalled replication forks. For example, loss of ZRANB3, HLTF, or SMARCAL1 abolishes formation of the reversed fork structures targeted by the MRE11 nuclease and thus results in chemoresistance of BRCA2-deficient cells (16). Inhibition of MRE11 (13,14), or loss of PTIP (which recruits MRE11 to reversed forks) (36), or of RADX (which inhibits RAD51 accumulation to stalled replication forks) (38) can similarly rescue PARPi sensitivity of BRCA2-deficient cells without restoring HR. Finally, inhibition of a parallel fork degradation pathway governed by the chromatin modifier EZH2 which recruits the nuclease MUS81 to stalled forks (39), or channeling the processing stalled forks towards translesion synthesis-mediated lesion bypass rather than fork reversal (35), can also suppress chemosensitivity of BRCA2-deficient cells. Here, we show that loss of E2F7, a transcriptional suppressor of RAD51, also results in protection against degradation of stalled replication forks. Our data suggest that increased RAD51 levels are enough to promote its loading to stalled replication forks even in the absence of BRCA2 activity.

Surprisingly, we found that E2F7 knockdown also restores HR in BRCA2-deficient cells. This effect is also mediated through RAD51 expression, as E2F7 knockdown could not rescue the HR defect of RAD51-depleted cells, thus ruling out an involvement of other E2F7 targets involved in HR (such as BRCA1). Moreover, this is not due to an indirect effect of E2F7 depletion on cell cycle distribution, as we did not observe any difference between the cell cycle profiles of BRCA2-depleted or BRCA2/E2F7-codepleted cells (not shown). How RAD51 is loaded to DSB ends in BRCA2-depleted cells is not clear, but it is likely to involve the activity of RAD52 which has been implicated in RAD51 loading in BRCA2-deficient cells (40).

At this time, it is unclear what is the relative contribution of HR rescue and fork protection activities to the E2F7-mediated chemoresistance of BRCA2-depleted cells. Nevertheless, to our knowledge this is the first time that restoration of HR (independent of a reversion mutation) is identified as a mechanism of PARPi resistance in BRCA2-deficient cells. Recently, E2F7 was reported to bind to double strand break sites and inhibit their repair, potentially through altering chromatin status at these sites (41). While it is not known if this activity requires RAD51, it is nevertheless possible that the rescue of BRCA2 chemosensitivity by loss of E2F7 also reflects this transcription-independent role of E2F7 in repressing DNA double strand break repair.

## ACKNOWLEDGEMENT

We would like to thank Dr. Jeremy Stark for the U2OS DR-GFP cells, Dr. Mark Stahl for the SH-SY5Y cells, Dr. Raymond Hohl for the HCC1395 cells, and the Penn State College of Medicine Flow Cytometry and Imaging cores.

